# A CDCA2 - MYC positive feedback loop controls cancer cells survival

**DOI:** 10.1101/2025.02.28.640789

**Authors:** Konstantinos Stamatiou, Lorena Ligammari, Malki Bothota, Antonia Tsavou, Ezgi Gokan, Ylenia Cicirò, Mariano F. Zacarías-Fluck, Laura Soucek, Arturo Sala, Paola Vagnarelli

## Abstract

MYC transcription factors are encoded by a small family of genes which include, in addition to the prototype member *MYC*, *MYCN* and *MYCL*. The majority of human cancers display expression alterations of *MYC* genes and their products, making this group of oncoproteins among the most relevant therapeutic targets in oncology. MYC is regulated at multiple levels, and protein phosphorylation is a critical mechanism used by the cell to modulate MYC stability and activity. Although there is a reasonable knowledge of the kinases required for MYC modifications, the counteracting phosphatases have been relatively understudied. Here we have investigated the role of the chromatin remodelling factor and protein phosphatase 1 (PP1) regulatory subunit CDCA2, also known as RepoMan, in the regulation of MYC proteins in cancer cells. Using RNA interference and regulatable degron-mediated degradation of CDCA2, we demonstrate that the PP1 subunit is required for MYC and MYCN stabilisation and viability of triple negative breast cancer, neuroblastoma and colon cancer cells. Proximity ligation assays indicate that both MYC and MYCN are in close proximity of CDCA2 *in vivo*. Furthermore, chromatin immunoprecipitation data and promoter mutation studies establish that *CDC2A* is a *bone-fide* MYC target gene in cancer cells, revealing a reciprocal regulatory loop that could be exploited for therapeutic purposes.

## INTRODUCTION

MYC is a transcription factor that regulates diverse cellular functions including cell proliferation, apoptosis, pluripotency maintenance, cellular reprogramming, and differentiation (Zacarias-Fluck *et al*, 2024). The MYC family encompasses the prototype member MYC and two other members, MYCN and MYCL, which encode oncoproteins sharing a similar structure, including a transcription activation domain at the amino terminus and a basic Helix Loop Helix-leucine zipper (bHLHZ) domain at the carboxyl terminus involved in DNA binding(Diolaiti *et al*, 2015). The expression of MYC family genes is altered in the majority of human cancers, largely as a consequence of gene rearrangement or amplification, and is a driver of aggressive behaviour of tumour cells via target gene-dependent or independent transcription(Baluapuri *et al*, 2020),(Jakobsen & Siersbaek, 2025).

MYC proteins regulate gene transcription in association with MAX(Sabo & Amati, 2014). The MYC-MAX heterodimer binds to E-box and non-E-box containing regulatory regions of 10–15% of all mammalian genes (Eilers & Eisenman, 2008). MYC is also a short half-life protein (∼30 min), that is primarily regulated by the GSK3/SCF/FBXW7 pathway. Mitogen regulated kinases phosphorylate MYC at serine 62 (S62) and GSK3β then phosphorylates threonine 58 (T58), which triggers protein phosphatase 2A (PP2A)-mediated S62 dephosphorylation. This recruits the SCFFBXW7 E3 ligase to direct MYC ubiquitylation and subsequent proteasomal degradation (Boi *et al*, 2023).

However, MYC can be phosphorylated at many other sites and these phosphorylations are important for MYC stability or function and several kinases have already been identified as responsible for modifications of specific sites. For example, Aurora B kinase phosphorylates MYC at serine 67 and promotes its protein stability(Jiang *et al*, 2020), CDK2 can phosphorylate S62 and prevents Ras-induced senescence(Hydbring *et al*, 2010; Hydbring & Larsson, 2010), while the stress response kinase Pak2 phosphorylates T358, S373, and T400 to reduce MYC activity, as it does phosphorylation by PKCζ at S373 (Kim *et al*, 2013).

The counteracting phosphatases for all these sites are so far unknown but recent work suggested that Protein phosphatase 1 (PP1) is a key phosphatase for MYC as inhibition of the catalytic subunit PP1c (or RNAi of PP1c) leads to increased phosphorylation levels of several MYC phosphosites (Dingar *et al*, 2018).

PP1c in cells only exists in complex with other proteins that are Regulatory Interactors of Protein Phosphate 1 (RIPPOs). They target PP1 to substrates or modulate its catalytic activity and more than 200 RIPPO/PP1 complexes have been identified so far. Therefore, searching for the relevant MYC – PP1 phosphatase complex is an important and challenging quest.

So far, only PNUTS/PP1 has been identified as a MYC phosphatase complex on chromatin(Dingar *et al*., 2018; Wei *et al*, 2022) but the specificity for the sites still remains obscure. However, several others PP1/RIPPO complexes are known to modulate chromatin states and dynamics and many have been associated with cancer. Here we have discovered that another RIPPO, CDCA2 (also known as Repo-Man), interacts with MYC and it is important for both MYC and MYCN stability. We have also identified *CDCA2* expression as a very powerful prognostic marker for Triple Negative Breast Cancer (TNBC) and *MYCN* amplified neuroblastomas.

## RESULTS

### *CDCA2* is co-expressed with *MYC* in Triple Negative breast cancer and is essential for cancer cell survival

Although PP1 has been implicated in MYC regulation, the mechanisms are not well understood. In several cell lines, PP1 inhibition by Caliculin A or PP1 knock-down, significantly increased MYC phosphorylation leading to its degradation(Dingar *et al*., 2018). To understand the role of PP1 in MYC biology, it is important to identify the RIPPO(s) that mediate this function. Recent work has shown that PNUTS (PPP1R10) is one of the effectors of c-MYC stability on chromatin (Dingar *et al*., 2018) (Finkelman *et al*, 2023; Wei *et al*., 2022).

However, considering the multiplicity of phosphosites within the MYC protein (Dingar *et al*., 2018), and the different types of chromatin it binds to (activator / repressor) it is plausible that multiple RIPPOs/PP1 complexes could be involved in its regulations at distinct types of chromatin(Jakobsen & Siersbaek, 2025; Lashen *et al*, 2023).

We therefore conducted an initial screening using the USCS cancer browser Xena (https://xena.ucsc.edu/welcome-to-ucsc-xena/) and the TCGA dataset for breast cancer to identify chromatin-associated RIPPOs linked to TNBC (a MYC driven cancer) (Fallah *et al*, 2017).

We selected NIPP1 (PPP1R8), PNUTS (PPP1R10), CDCA2 (PPP1R81) RIF1 and MKI-67 as all these RIPPOs have been associated with cancer before (Dingar *et al*., 2018),(Baxter *et al*, 2024; Egger *et al*, 2016; Gu *et al*, 2022; Harvey-Jones *et al*, 2024; Li *et al*, 2023; Marx *et al*, 2020; Peng *et al*, 2010; Sun *et al*, 2023; Tang *et al*, 2021; Wei *et al*., 2022; Zhou *et al*, 2024).

When comparing their mRNA levels with *MYC* expression, the analyses clearly highlighted a strong correlation for *PPP1R81(CDCA2*) and *MYC*, even stronger that the one with *Ki-67*, a proliferation marker used in pathology to stage breast cancer (Finkelman *et al*., 2023). CDCA2 is also known as Repo-Man, a chromatin associated RIPPO important for cell division (Peng *et al*., 2010) (de Castro *et al*, 2017) (de Castro *et al*., 2017; Qian *et al*, 2013; Qian *et al*, 2011; Trinkle-Mulcahy *et al*, 2006; Vagnarelli, 2014; Vagnarelli *et al*, 2006; Vagnarelli *et al*, 2011) (**Figure EV1 A**). *CDCA2* mRNA levels correlate with the stage of breast cancer (**Figure 1 A**) where higher expression is linked to more advanced stages, and they negatively impact on the overall survival of breast cancer cases (**Figure 1 B**).

**Figure 1.**
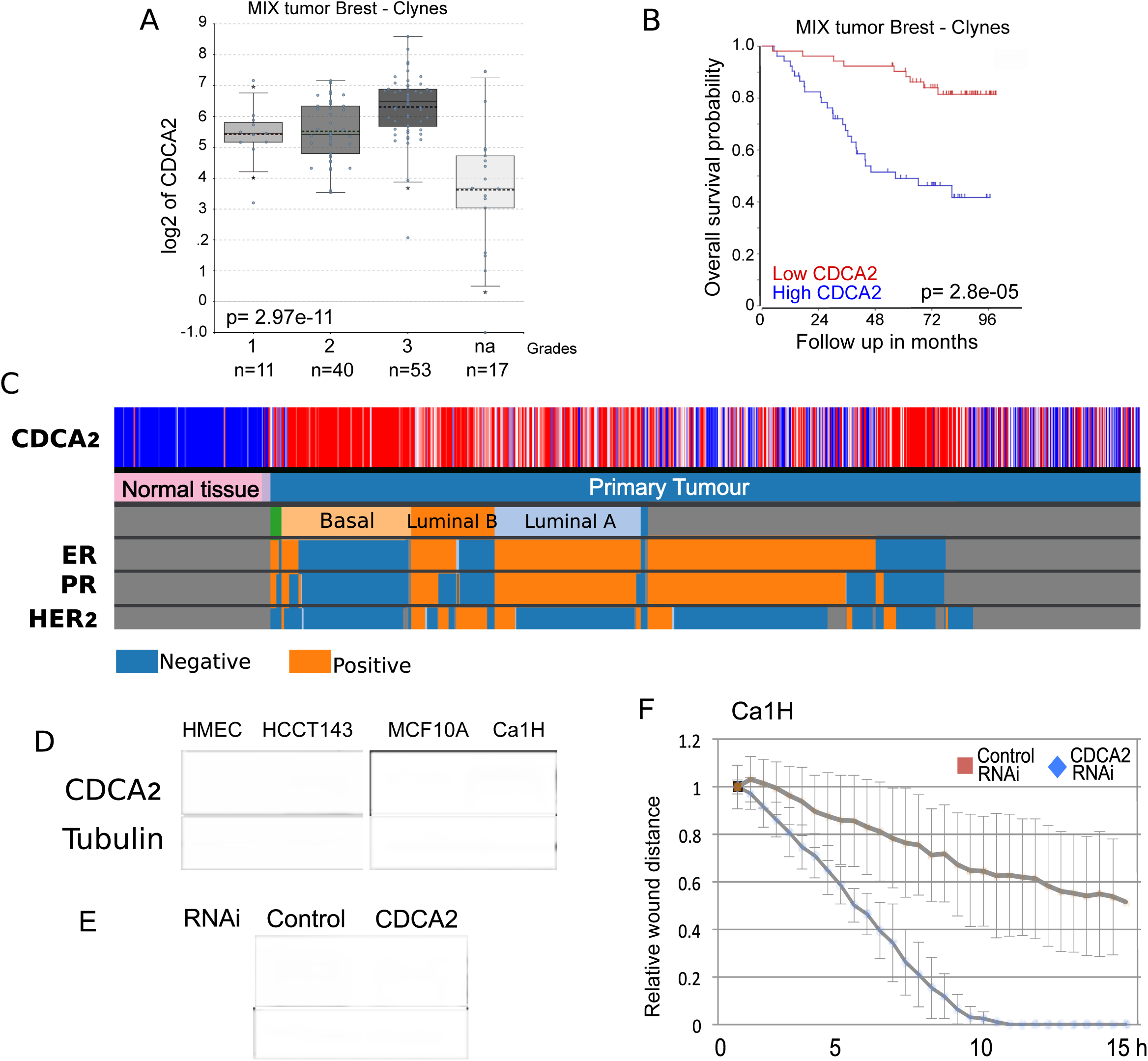
**A)** CDCA2 expression levels in mix breast tumours (Clynes dataset) of different grades. **B)** Kaplan–Meier curve (survival probability in months) of breast tumor patients expressing low (red curve) or high (blue curve) levels of CDCA2. **C)** UCSC cancer browser analyses of CDCA2 levels in Breast cancer normal breast and tumours stratified for Oestrogen receptor (ER), Progesteron (PR), Her 2 expression and type. Blue represents low expression and red high expression levels. **D)** Western blot of HMEC, HCCT143, MCF10A and Ca1H cell lines. The blots were probed with anti-alpha tubulin and with anti-CDCA2 antibodies. **E)** Representative Western blot of Ca1H cell line transfected with control-Si or CDCA2-Si oligos for 36 h. The blot was probed with anti-alpha tubulin and with anti-CDCA2 antibodies. **F)** Quantification of relative wound distance of Ca1H cell line transfected with control Si (orange) or CDCA2 Si (blue) oligos. The values represent the average of 3 independent replicas, and the error bars are the standard deviations. The experiments were analysed by Student’s t-test.***p < 0.001

Interestingly, when stratified for cancer subtypes, *CDCA2* expression levels were significantly higher in TNBC (**Figure 1 C and Figure EV1 A**). The higher mRNA expression also correlates with higher protein levels in a TNBC cell line (HCCT143) and in the isogenic invasive MCF10A derivative Ca1H (also known to have high MYC levels (Tang *et al*., 2021)) compared to normal mammary epithelial (HMEC) and the non-transformed MCF10A cells (**Figure 1 D**). Altogether these data suggest a positive correlation between *MYC* expression and *CDCA2* that we sought to explore further.

We next investigated if CDCA2 was essential for cell survival in these cell types by a SiRNA approach using the Ca1H cell line. We could successfully knock-down CDAC2 in the Ca1H cells (**Figure 1 E**) and the depletion caused cell death as shown by the significant increase of sub-G1 cells (**Figure EV1 B**) within 48h. CDCA2 was also essential for wound closure in wound-healing assays (**Figure 1 F**).

CDCA2 is a chromatin bound protein. We previously mapped its preferred binding sites on HeLa cells chromatin and showed that CDCA2 is important for the repression of several PRC2 regulated genes (de Castro *et al*., 2017). We therefore checked if CDCA2, which is overexpressed in TNBC, could also act as a repressor in this context. To this purpose, we analysed the gene expression profiles of the previously identified CDCA2 bound genes in normal mammary epithelium and TNBC samples using the USCS cancer genomic platform. The analyses showed that the majority of CDCA2 bound genes negatively correlate with CDCA2 expression in this cancer subtype and that the highest represented category belongs to the GO terms of cell adhesion components (**Figure EV1 C**). Interestingly, 69.2% of them harbour a MYC binding site at their promoter regions.

These analyses altogether let us to conclude that CDCA2 is an important regulator of gene expression in TNBC and it is essential for cell viability.

### CDCA2 phosphatase activity and chromatin binding are essential for its function in TNBC

CDCA2 is a protein with several domains. These include a) a region binding to PP1 and b) a binding site for chromatin (**Figure 2 A**). To test if these functions were necessary for wound healing in TNBC, we performed a set of rescue experiments with oligo-resistant GFP-tagged mutant forms of the protein in a CDCA2 RNAi background. The experiments were conducted in the TNBC cell line HCTT143 where, similarly to what we obtained for the Ca1H cell line, CDCA2 RNAi led to a defect in the closure of the wound (**Figure 2 B**). In these settings, while the GFP-tagged *wt* form of CDCA2 was able to rescue the wound closure defect caused by CDCA2 RNAi, neither the chromatin binding mutant (GFP-CDCA2*^S893D^*) nor the one unable to bind PP1 (GFP-CDCA2*^RAXA^*) or GFP alone were able to do so (**Figure 2 C-F**). These experiments confirm that CDCA2 function in TNBC is PP1-dependent and requires binding to chromatin.

**Figure 2.**
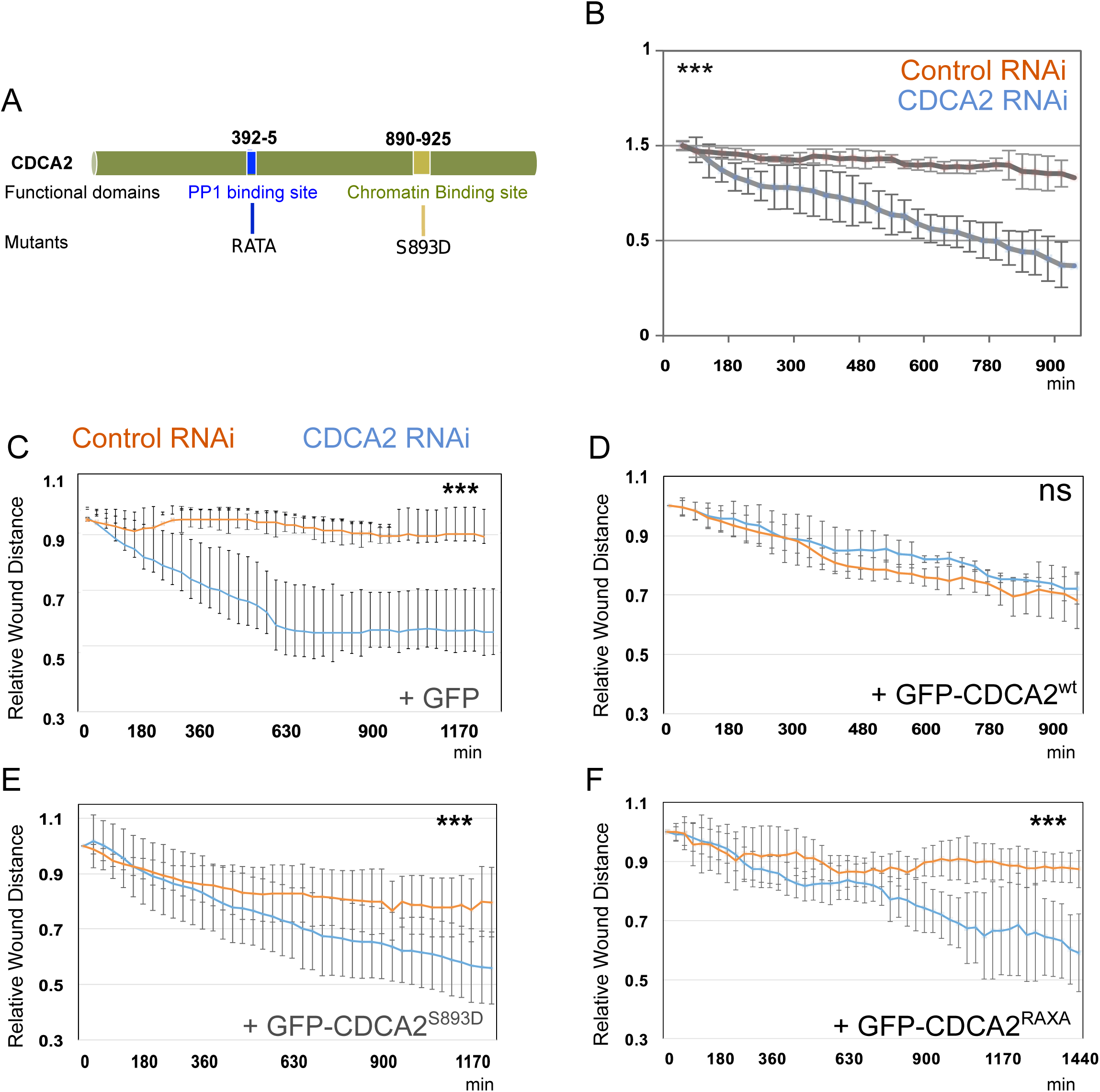
**A)** Schematic diagram of CDCA2 and its domains. RATA mutation at the PP1 binding site 392-6 and S893D mutation at the chromatin binding site 890-925 amino acids are highlighted in Blue and light green respectively. **B)** Quantification of the relative wound distance of HCCT143 cells transfected with control Si (orange) or CDCA2 Si (blue) oligos for 36 h. The values represent the average of 3 independent replicas, and the error bars are the standard deviations. The experiments were analysed by Student’s t-test.***p < 0.001 **C-F)** Quantification of relative wound distance of cells co-transfected with control Si (orange) or CDCA2 Si (blue) oligos and GFP (C), or oligo-resistant GFP-CDC2A^wt^ (D), GFP-CDC2A^S893D^ (E) and GFP-CDC2A^RAXA^ (F) plasmids for 36 h. The values represent the average of at least 3 independent replicas, and the error bars are the standard deviations. The experiments were analysed by Student’s t-test. ns=no significant, ***p < 0.001

### *CDCA2* is a MYC regulated gene

Because of the strong correlation between *CDCA2* and *MYC* expression in TNBC, we wondered if *CDCA2* could be itself a MYC-regulated gene. Further inspection of the *CDCA2* promoter revealed the presence of three E-box MYC binding sites (Gaubatz *et al*, 1994), and MYC ChIP datasets, showed in both MCF-7 and HeLa cells that MYC is present at the *CDCA2* promoter (**Figure 3 A**).

**Figure 3.**
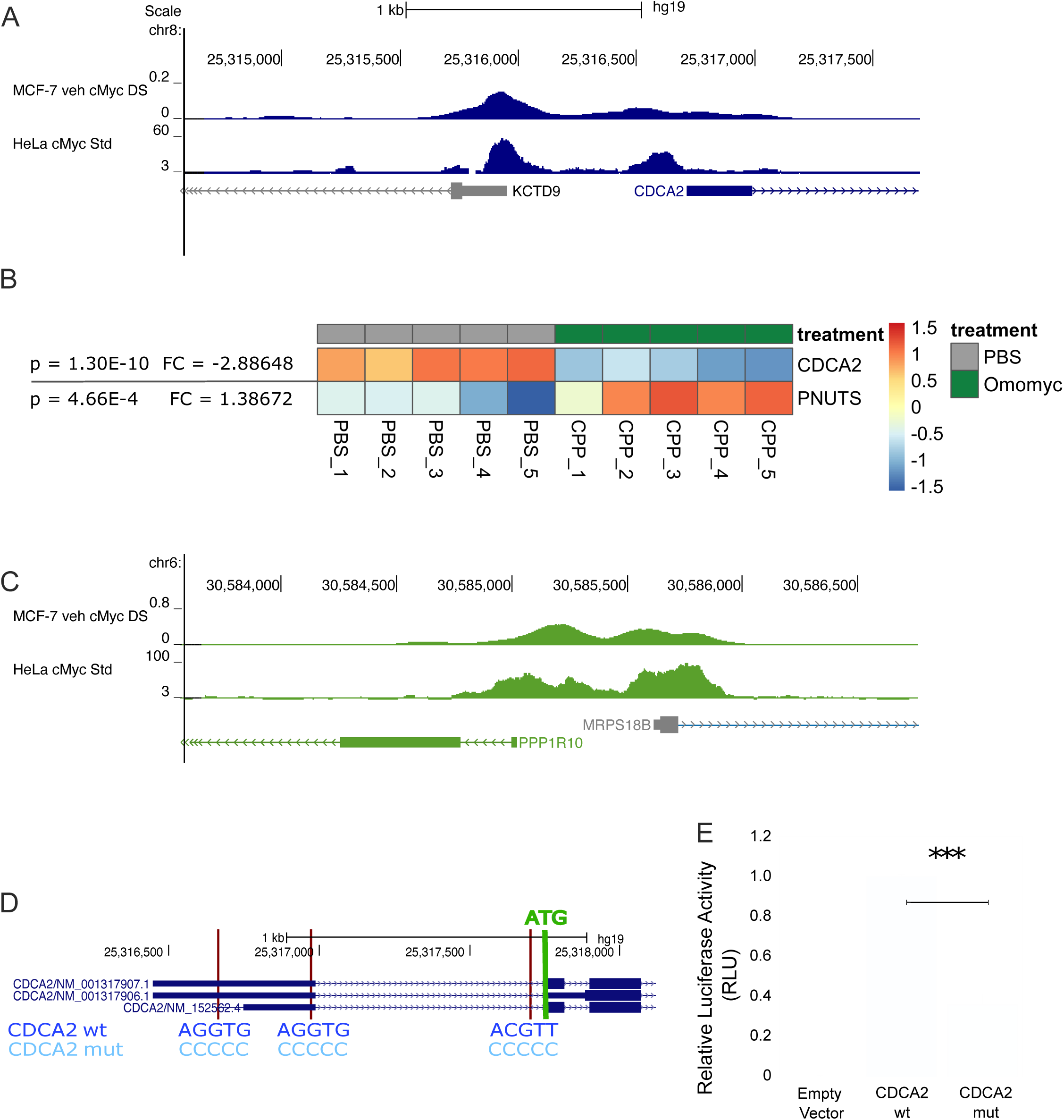
**A)** cMYC chromatin immunoprecipitation sequencing profiles at the CDCA2 locus in MCF-7 and HeLa cell lines **B)** cMYC chromatin immunoprecipitation sequencing profiles at the PNUTS (PPP1R10) locus in MCF-7 and HELA cell line.s **C)** Microarray analysis MDA-MB231 cells treated with 20 µM Omomyc (CPP) or PBS for 3 days. P-values, fold change (FC) and heatmaps for CDCA2 and PNUTS. **D)** Schematic representation of the genomic locations selected to be mutated for the pGL3-CDCA2 wt/mut constructs used for the luciferase assay. The core MYC (AG/CGTG/T) binding sequence and the mutations introduced are indicated by the red lines at the promoter region of the CDCA2 gene. **E)** The bar plot indicates the RLU (Relative Light Unit) values corrected for transfection efficiency with pRenilla luciferase. pGL3 empty vector was used to calculate the background luciferase activity. The values represent the average of 3 independent replicas and 3 technical replicas, and the error bars are the standard deviations. The experiments were analysed by Student’s t-test. ns=no significant, ***p < 0.001

MYC inhibition can be achieved by a mini-protein called omomyc. Omomyc is a dominant-negative homologous to the bHLHZ of MYC, containing point mutations that allow homo- and hetero-dimerisation with MAX and all MYC-family proteins(Masso-Valles & Soucek, 2020). Omomyc has been recently validated as an anti-cancer agent in patients with solid tumours in a Phase I trial (Garralda *et al*, 2024). Using this mini-protein inhibitor in the TNBC cell line MDA-MB231, *CDCA2* expression decreased by 2.9 fold (**Figure 3 B**), thus suggesting a link between MYC and *CDCA2* expression.

As MYC has also been linked to another RIPPO, PNUTS, whose promoter also harbours MYC binding sites as well (**Figure 3 C**), we checked if this PP1 interactor was also MYC regulated. However, to our surprise, in this setting, Omomyc expression led to an increase in *PNUTS* mRNA of 1.4 folds, the opposite effect of *CDCA2* (**Figure 3 B**). This may suggest that MYC acts more as a suppressor for PNUTS at least in this cell type. Indeed, public TNBC datasets indicate that *PNUTS* (*PPP1R10*) expression is negatively correlated with that of MYC (**Figure EV1 A** – cnfPPP1R10 and MYC).

We then wanted to confirm that *CDCA2* was indeed a MYC target. For this, we used a luciferase reporter assay where the *CDCA2* promoter containing the E-boxes was used to drive the expression of luciferase. Transfection of the *CDCA2^w^*^t^ promoter in the Ca1H cell line could drive the luciferase expression however, when all the E-boxes of the *CDCA2* gene were mutated, the promoter activity was significantly decreased (**Figure 3 D, E**). This clearly shows that *CDCA2* is a MYC target gene and that it is essential for proliferation of MYC-driven breast cancer cells.

### CDCA2 is required for c-MYC stability

We then wondered how CDCA2 activity in TNBC was linked to MYC biology. MYC stability/degradation is regulated by phosphorylation and the wound healing experiments in breast cancer cells indicated that CDCA2 function is PP1-mediated. We therefore depleted CDCA2 in the Ca1H and the HCCT143 cell lines and quantified MYC levels using immunofluorescence. The results clearly show that a significant decrease in MYC levels was achieved upon CDCA2 depletion in both cell lines (**Figure 4 A, B**).

**Figure 4.**
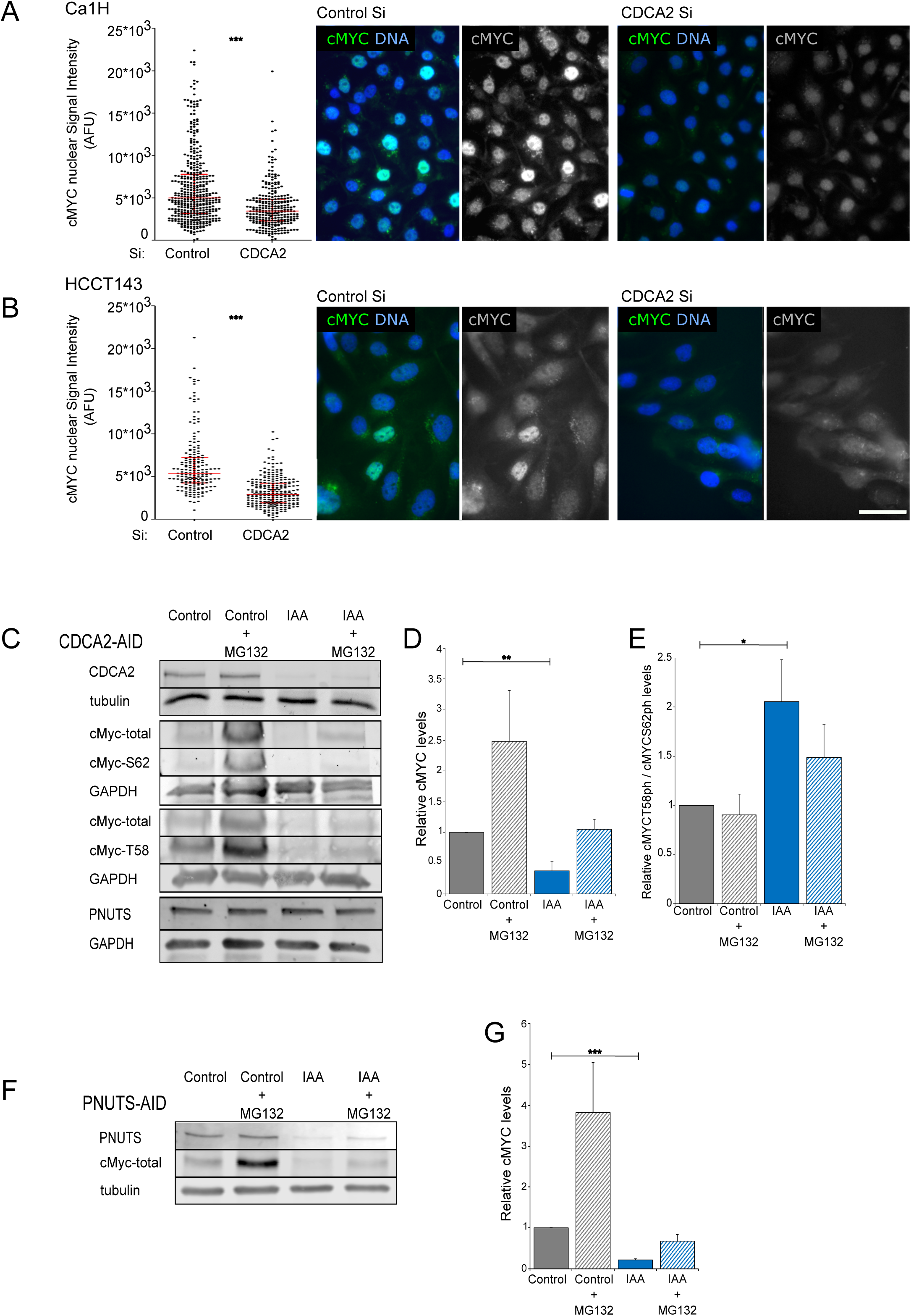
**A and B)** Violin plot of the quantification of cMYC nuclear signal intensity (left). Whiskers are the upper and lower adjacent values and the line is the median. A Wilcoxon test was conducted for comparing the experiments. ***p < 0.001 (Left) Representative images of the Ca1H (A) and HCCT143 (B) cell lines 48 hours post transfection with control or CDCA2 oligos, stained with cMYC antibodies (green) and counterstained with DAPI (blue). **C)** Representative Western blot analyses of HCT116-CDCA2-AID cell line treated with doxycycline (2 μg/ml) for 24 h before the addition of IAA, then untreated (Control) or treated with IAA (IAA) for 24 h followed by a 2 h treatment with 20μM MG132 in the presence (IAA + MG132) or absence Control + MG132) of IAA. The blots were probed with anti-GAPDH or anti-alpha tubulin antibodies and with anti-CDCA2, anti-PNUTS, anti-cMYC-total, anti-cMYC-S62ph and anti-cMYC-T58ph antibodies. The images were acquired with a LICOR Imaging system in the linear range for quantification purposes. **D)** Quantification of total cMYC from the experiments in (C). The values represent the average of 3 independent replicas, and the error bars are the standard deviations. The experiments were analysed by Student’s t-test. **p < 0.01. **E)** Quantification of cMYC-T58ph / cMYC-S62ph from the experiments in (C). The values represent the average of 3 independent replicas, and the error bars are the standard deviations. The experiments were analysed by Student’s t-test. *p < 0.05. **F)** Representative Western blot analyses of HCT116-PNUTS-AID cell line treated with doxycycline (2 μg/ml) 24 h before the addition of IAA, then untreated (Control) or treated with IAA (IAA) for 24 h followed by a 2 h treatment with 20μM MG132 in the presence (IAA + MG132) or absence (Control + MG132) of IAA. The blots were probed with anti-alpha tubulin, anti-PNUTS and anti-cMYC-total antibodies. The images were acquired with a LICOR Imaging system in the linear range for quantification purposes. **G)** Quantification of total cMYC from the experiments in (C). The values represent the average of 3 independent replicas, and the error bars are the standard deviations. The experiments were analysed by Student’s t-test.***p < 0.001

As RNAi could exert off-target effects (although we have shown that the wound-healing can be rescued by GFP-CDCA2^wt^), we took advantage of a more accurate system we recently developed to deplete CDCA2 using the endogenously tagged CDCA2 allele with an auxin degron module (AID) where the OSTR1 component is inducible by Doxycycline (DOX). Addition of IAA to the system triggers the degradation of the specific protein, thus avoiding any possible off-target effect that the RNAi approach could cause (the validation of the cell line is in (Stamatiou *et al*, 2024)). This cell line has been generated in the HCT116, a colorectal cancer cell line in which MYC expression is elevated and essential for survival (Lu *et al*, 2010).

Using this additional system, we could show that CDCA2 degradation leads to a decrease of MYC protein levels (**Figure 4 C, D**). These data strengthen again the relationship between MYC and CDCA2 and indicate that whilst MYC is important for *CDCA2* expression, CDCA2 is also important for maintaining MYC levels.

As previous work has linked the stability of MYC protein levels to PNUTS (Dingar *et al*., 2018), we wondered if the phenotype we observed was caused by an effect of CDCA2 on PNUTS. However, our analyses clearly show that CDCA2 degradation does not change PNUTS protein levels in HCT116 (**Figure 4 C**). To verify that the reported stabilization of MYC by PNUTS was also occurring in HCT116 and was specific (previous data were only conducted by PP1 knock down and inhibition), we used a similar cell line where we have endogenously tagged PNUTS with the same degron module as CDCA2 (Stamatiou *et al*., 2024). Here, we could confirm that, upon addition of IAA and PNUTS degradation, MYC protein levels decrease as previously reported (**Figure 4 F, G**) (Dingar *et al*., 2018). Overall, these data indicate that there are at least 2 RIPPOs involved in MYC regulation.

As MYC stability is regulated by its phosphorylation status, we investigated the levels of the T58ph and S62ph sites. The western blot analyses revealed that, upon CDCA2 degradation, the ratio between the T58ph and S62ph levels is increased in the presence of IAA, a signature that favours MYC degradation (**Figure 4 C, E**).

The decrease in MYC protein levels could be due to either increased degradation or a arrest in G1 at the restriction checkpoint caused by the depletion of CDCA2 or PNUTS. In fact, previous work in Human Squamous Cell Carcinoma cancer cells have shown a G1 arrest upon *CDCA2* SiRNA (Uchida *et al*, 2013), and recent work from our lab has highlighted a G1 block in cells where either CDCA2 or PNUTS were degraded in mitosis and then released (Stamatiou *et al*., 2024). We therefore set out to test these possibilities using the AID tagged cell lines for PNUTS and CDCA2. In these experiments, we arrested the cells in thymidine (after the commitment stage and when MYC is already expressed), then we degraded the proteins for 4 h and analysed MYC phosphorylation status and gene transcription levels.

At this cell cycle stage, degradation of either CDCA2 or PNUTS lead to a decrease of MYC protein levels and changes in the phosphorylation balance (**Figure EV2 A-C, D-F**). As previously seen for the asynchronous population, the ratio between MYC T58ph/S62ph increases upon degradation of each protein, a signature for increased degradation. These experiments ruled out the hypothesis of the effect being the consequence of cell cycle arrest in early G1.

CDCA2 depletion does not cause significant changes in either *CDCA2* or *PNUTS* mRNA. To further analyse the relationship between the two RIPPOs, we also investigated their relative protein levels in the degradation experiments. While CDCA2 degradation does not affect the protein levels of PNUTS (**Figure 4 C**), degradation of PNUTS affects CDCA2 protein levels (**Figure EV2 G-H**) but not its transcription (**Figure EV2 I**). However, either PNUTS or CDCA2 degradation decreases *MYC* expression at the G1/S transition (**Figure EV2 I, J**). Interestingly, degradation of PNUTS, increased its own transcription (**Figure EV2 J**).

These data altogether suggest that both RIPPOs are important for MYC regulation and stability, and they seem to place PNUTS upstream of CDCA2, as degradation of PNUTS does affect CDCA2 levels.

Interestingly, we have also discovered a feedback loop in PNUTS regulation whereby its degradation leads to increase in expression (**Figure EV2 J**).

### CDCA2 is regulated by MYCN in neuroblastoma and is essential for MYCN stability

Previous studies have shown that *CDCA2* is classified into a unique group of genes up-regulated during the progression of neuroblastomas (Krasnoselsky *et al*, 2005). Since these tumours are often associated with *MYCN* amplifications, we wondered if CDCA2 could also be linked to MYCN in this pathological condition.

We therefore first conducted in silico analyses of neuroblastoma gene expression datasets using the R2 genomic platform and interrogated the relationship between *MYCN* expression and the same set of chromatin-associated PP1 phosphatases used before for breast cancer. The analyses revealed that among all the RIPPOs investigated, *CDCA2* was the one where the expression levels were positively correlated with *MYCN* amplification and metastatic stage 4 (**Figure 5 C, D)** but also that expression of CDCA2, but not of other RIPPOs, was significantly associated with low patient survival (**Figure 5 F** and **Figure EV3 A-C**). This is in contrast with the expression levels of *PNUTS*, that has been previously linked to MYC regulation (**Figure 5 A, B, E**).

**Figure 5.**
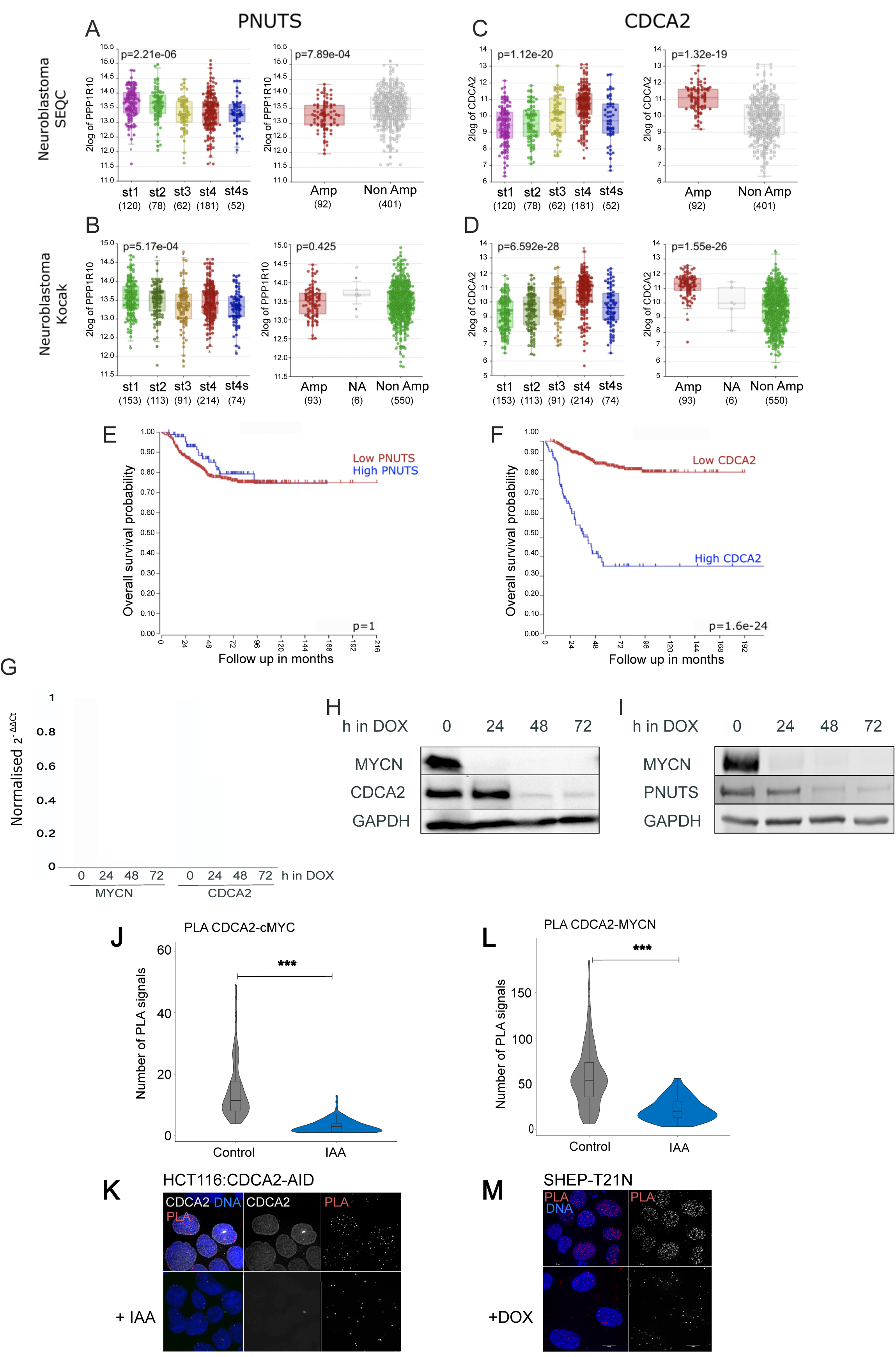
**A, B)** PNUTS expression levels in neuroblastoma patients with different stages (st) of cancer (left) and PNUTS expression levels in neuroblastoma patients with (red) and without (grey) MYCN amplification (right) (SEQC cohort -A, and Kocak cohort-B), the numbers indicate the sample sizes. **C, D)** CDCA2 expression levels in neuroblastoma patients with different stages (st) of cancer (left) and CDCA2 expression levels in neuroblastoma patients with (red) and without (grey) MYCN amplification (right) (SEQC cohort -C, and Kocak cohort-D), the numbers indicate the sample sizes. **E, F)** Kaplan–Meier curve (survival probability in months) of neuroblastoma patients expressing low (red curve) or high (blue curve) levels of PNUTS (E) or CDCA2 (F). **G)** qPCR analyses of MYCN and CDCA2 expression in SHEP-T21N treated without (0 h) or 1 μg/ml doxycycline for 24, 48 and 72 h. The data represent the average of 3 independent replicas and the error bars are the standard deviations. The experiments were analysed by Student’s t-test. *p < 0.05, ***p < 0.001 **H)** Representative Western blot analyses of SHEP-T21N treated without (0 h) or with 1 μg/ml doxycycline for 24, 48 and 72 h. The blots were probed with anti-alpha tubulin, anti-CDCA2 and anti-MYCN-total antibodies. **I)** Representative Western blot analyses of SHEP-T21N treated without (0 h) or with doxycycline for 24, 48 and 72 h. The blots were probed with anti-alpha tubulin, anti-PNUTS and anti-MYCN-total antibodies. **J)** Quantification of the experiment in (B). The box inside the violin represents the 75th and 25th percentile, whiskers are the upper and lower adjacent values and the line is the median. Sample size: control = 122, IAA = 126. A Wilcoxon test was conducted for comparing the experiments and ***p < 0.001. **K)** Representative images of the proximity ligation assay (PLA) using anti-cMYC and anti-GFP antibodies on HCT116:CDCA2-AID cell line with (top) or without (bottom) of IAA. **L)** Quantification of the experiment in (D). The box inside the violin represents the 75th and 25th percentile, whiskers are the upper and lower adjacent values and the line is the median. Sample size: control = 179, IAA = 123. A Wilcoxon test was conducted for comparing the experiments and ***p < 0.001 **M)** Representative images of the proximity ligation assay (PLA) using anti-cMYC and anti-CDCA2 antibodies on SHEP-T21N cell line with (top) or without (bottom) of doxycycline. Scale bar 5 μm.

Based on these findings and considering the knowledge we acquired on MYC and CDCA2 regulation in TNBC and colon cancer cells, we wondered if a similar relationship between CDCA2 and MYCN also occurred in neuroblastoma.

We first tested whether MYCN was bound to the *CDCA2* promoter in neuroblastoma cell lines. Cystrome analyses of MYCN ChIP-seq dataset revealed that MYCN was indeed present at the CDCA2 promoter in both SHEP-21N, engineered to conditionally express MYCN, and the naturally *MYCN* amplified Kelly cell line (**Figure EV3 D**), suggesting that MYCN could be important for *CDCA2* transcription. To verify this, we used a cell line that does not expresses endogenous levels of *MYCN* but carries a MYCN transgene under the control of a DOX repressible promoter (Lutz *et al*, 1996). Without DOX, MYCN is expressed, and the protein accumulates at high levels; upon addition of DOX, *MYCN* transcription is highly reduced after 24h (**Figure 5 G**) and MYCN protein levels are negligible in the cells (**Figure 5 H**). The analyses of CDCA2 in these settings showed that, without DOX, CDCA2 is highly expressed however, upon addition of DOX, *CDCA2* mRNA expression decreases as do its protein levels with a lag time compared to MYCN (**Figure 6 G, H**); this is expected as CDCA2 requires the passage through mitosis to complete its degradation (Manzione *et al*, 2020). Furthermore, we analysed the levels of CDCA2 in a few MYCN expressing cell lines and correlate their relative levels. The data showed that there is also a positive correlation between the levels of two proteins (**Figure EV3 F, G**).

Interestingly, *PNUTS* also shows accumulation of MYCN upstream of its TSS (**Figure EV3 E**), and repression of MYCN by addition of DOX in the SHEP-21 decreases PNUTS protein levels (**Figure 5 I**), although this does not seem to correlate with *MYCN* mRNA levels in neuroblastomas (**Figure 5 A, B**). Altogether these data indicate that also in neuroblastoma *CDCA2* is a MYCN regulated gene.

We therefore set out to investigate if MYCN stability in neuroblastoma was also linked to CDCA2 levels, as previously observed for MYC in breast cancer. We selected the MYCN amplified LAN-1 and SK-N-B(2)-C and the SHEP-T21N cell lines. In these cells we conducted *CDCA2* RNAi experiments and evaluated MYCN levels and its phosphorylation status. In all the cell lines, CDCA2 depletion leads to a decrease in MYCN due to increased degradation (as addition of MG132 restores MYCN levels) (**Figure EV4 A,B,D,E,G,H**). In addition, similar to what we observed for MYC, the T58ph/S62ph ratio was more elevated (**Figure EV4 C, F, I**), congruent with the observed MYCN degradation.

### CDCA2 interacts with MYC and MYCN

CDCA2 was previously identified as a MYC interacting protein in U2OS cells using an APEX-MYC proximity labelling approach (Solvie *et al*, 2022) and also in HEK293T cells by a MYC Bio-ID interactome, where it was shown to specifically interact with MYC homology boxes (MBs) MBIV (Kalkat *et al*, 2018).

However, these biochemical interactions have never been validated for MYC and no data are available for MYCN.

Because of the effect of CDCA2 depletion on MYC and MYCN stability, we wanted to test if CDCA2 is indeed a bona fide MYC or MYCN interactor in cells.

To this purpose, we conducted proximity ligation assays (PLA) and tested the interactions between the endogenous CDCA2 and MYC in the HCT116 cells and CDCA2 and MYCN in the TET-21 cell line. We could detect positive PLA signals confirming the interaction between CDCA2 and either MYC or MYCN, that were significantly decreased either upon CDCA2 degradation by IAA (**Figure 5, J K**) or MYCN repression by DOX (**Figure 5 L, M**).

## DISCUSSION

Despite many years of work and progress toward understanding MYC regulation, we are still far from establishing the specific contribution of its several post-translational modifications for MYC activity and stability. For example, MYC can be phosphorylated at many different sites but only very few have been linked to specific changes in MYC activity. Even less known are the phosphatases that reverse those modifications. The only established MYC phosphatase so far is PP2A that has been implicated in the de-phosphorylation of S62 and MYC degradation (Boi *et al*., 2023). A PP2A inhibitor, CIP2A, has been shown to be important for maintaining high MYC S62 phosphorylation, critical for MYC’s ability to re-initiate proliferation, and intestinal regeneration in response to DNA damage in mouse models (Myant *et al*, 2015). Interestingly, the major fraction of S62phMYC is bound to the Nuclear envelope via CIP2A and selectively supports the stability of the Lamina-associated pool of S62phMYC.

PP1 has been also implicated in MYC de-phosphorylation but the only RIPPO that has been so far characterised is PNUTS. PNUTS was identified in MYC-Bio-ID experiments, and it was shown that MYC and PP1/PNUTS can co-exist at promoters genome-wide. Inhibition or knock down of PP1 leads to the hyperphosphorylation of MYC and its dissociation from chromatin (Dingar *et al*., 2018). We and others have shown that PNUTS is a major regulator of transcription and that PNUTS degradation causes at least 2000 transcripts to be up or down-regulated (Stamatiou *et al*., 2024). PNUTS may control MYC phosphorylation at these highly transcribed loci. Although PNUTS physically interacts with MYC (Wei *et al*., 2022), PNUTS knockdown and rescue experiments have not been conducted so far. Moreover, it is not known if this link with MYC is also valid for MYCN.

MYC can both activate and repress genes. For example, three genes encoding cell-cycle inhibitory proteins, p15Ink4b, p21Cip1 and the MYC-antagonist Mad4, are repressed by MYC through interaction with Miz-1(van de Wetering *et al*, 2002). In addition, EZH2, the main enzymatic subunit of the polycomb repressive complex 2 (PRC2), directly interacts with the MYC family oncoproteins MYC and MYCN, and promotes their stabilization by competing against the SCF^FBW7^ ubiquitin ligase to bind MYC and MYCN (Wang *et al*, 2022). Furthermore, direct recruitment of EZH2 by MYCN is associated with repressive chromatin marks at the promoter of bivalent genes and their silencing (Corvetta *et al*, 2013).

PNUTS is not present at polycomb repressed genes, therefore it could be envisaged that other chromatin-bound phosphatases can also be involved in MYC regulation at different chromatin sites.

Here we have analysed a few chromatin-associated RIPPOs that have been linked to cancer and discovered that CDCA2 is another important phosphatase interacting with both MYC and MYCN. We have shown that depletion or degradation of CDCA2 in different cellular systems leads to decreased MYC levels and increased T58 phosphorylation. Interestingly, we have previously shown that CDCA2 binds H3K27me3 and is enriched at bivalent genes. This suggests that maybe different types of chromatin may need to recruit different PP1/MYC regulators.

In the future it will be important to understand which PP1-dependent phosphosites are specific for each RIPPO, or if there is no specificity but just compartmentalisation of the phosphatase activity to specific chromatin regions achieved by the different RIPPOs.

We have also shown that *CDCA2* is a MYC target gene and its expression is of prognostic value both for Triple Negative Breast cancers and for MYCN amplified neuroblastoma.

## Materials and methods

MCF10A, MCF10A-TK1 and MCF10A-CA1h were obtained from the Karmanos Cancer Institute (via MTA) and maintained in DMEM/F12 (Invitrogen) supplemented with: 10% foetal bovine serum (FBS) (Labtech), 1% Penicillin-Streptomycin (Gibco), 10 µg/ml Insulin (Sigma), 20 µg/ml EGF (Perprotech), 0.5 µg/ml Hydrocortisone (Sigma).

HCC1143 and SK-N-BE(2)-C cells were grown in Gibco RPMI 1640 GlutaMAX™ supplemented with 10% foetal bovine serum (FBS) (Labtech) and 1% penicillin– streptomycin (Gibco) at 37 °C with 5% CO2.

SHEP-T21N and LAN-1 cells were grown in Gibco DMEM GlutaMAX supplemented with 10% foetal bovine serum (FBS) (Labtech) and 1% penicillin–streptomycin (Gibco) at 37 °C with 5% CO2.

HCT116-CDCA2-AID and HCT116-PNUTS-AID cells were grown in Gibco™ McCoy’s 5A Medium GlutaMAX supplemented with 10% foetal bovine serum (FBS) (Labtech) and 1% penicillin–streptomycin (Gibco) at 37 °C with 5% CO2.

### Transfections

For siRNA treatments, HCCT143, Ca1H, SK-N-BE(2)-C, SHEP-T21N and LAN-1 cells were seeded into 6-well plates, transfected using Polyplus JetPrime® (PEQLAB) with the appropriate siRNA oligonucleotides (50 nM) and analysed after 30-48 h. The siRNAs oligoes were published before (Vagnarelli *et al*., 2011) and obtained from Merck.

### Immunofluorescence microscopy

Cells were fixed in 4% PFA and processed as previously described (Vagnarelli *et al*., 2006). Primary and secondary antibodies were used as listed in the table. Three-dimensional datasets were acquired using wide-field microscope (NIKON Ti-E super research Live Cell imaging system) with a 100X Plan Apochromat lens, numerical aperture (NA) 1.45.

The datasets were deconvolved with the NIS Elements AR analysis software (NIKON). Three-dimensional datasets were converted to maximum projection, exported as TIFF files, and imported into Inkscape for final presentation.

### Immunoblotting

Whole cell extracts were prepared by direct lysis in 1× Laemmli sample buffer (Laemmli, 1970).

Membranes were blotted with primary and secondary antibodies that are listed in the Table. Membranes were visualised using either the Bio-Rad ChemiDoc XRS system or the LiCor Odyssey system.

HMEC whole cell lysates were provided by Prof Newbold, Brunel University of London.

### Flow cytometry cell cycle analysis

Cells were trypsinised, resuspended and incubated at room temperature for 30 min in 70% ice-cold ethanol. Cells were centrifuged at 1000 g for 5 min and washed with PBS, and the supernatant was discarded. The pellet was resuspended in 200 μl of RNase A/PBS (100 μg/ml) and incubated for 2 h at 37 °C in the dark. Propidium iodide (Fisher Scientific, P3566) was added at a final concentration of 5 μg/ml just before analysing the samples by flow cytometry using the ACEA Novocyte Flow Cytometer. The analysis was performed using the NovoExpress® software.

### Proximity ligation assay (PLA)

Proximity ligation assay was performed according to the manufacturer’s protocol (Sigma). SHEP-T21N doxycycline (1 μg/ml) for 24 h and HCT116-CDCA2-AID cells were treated with doxycycline (1 μg/ml), 24 h before the addition of auxin 1000 μM for 4h. The cells were fixed, permeabilised and blocked with BSA as previously described (Vagnarelli *et al*., 2006).

The antibodies were used at the following concentrations: 1:100 anti-CDCA2, 1:100 anti-MycN, 1:100 anti-cMyc and 1:10,000 anti-GFP [PABG1] (Cat# PABG1-20, RRID:AB_2749857). PLA probes were added, and ligation was performed following manufacturer’s instructions (Sigma). Coverslips were mounted and observed on the previously mentioned wide-field NIKON microscope.

### Wound Healing Assay: live imaging and analysis

2 × 10^5^ cells/ well in 1 ml of medium were grown overnight in a 2-well chamber and transfected the following day with siRNA, oligo control or oligo against Repo-Man (50 nM) and 500ng of GFP or GFP-CDC2A^wt^ or GFP-CDC2A^S893D^ or GFP-CDC2A^RAXA^ plasmid for 36 h. After 24h the medium in each chamber was replaced with Leibnovitz’s phenol free medium, supplemented with antibiotics and serum. A wound was made in each well using a sterile tip and the cells were transferred on a wide-field microscope (NIKON Ti-E super research Live Cell imaging system) at 37°C. Several points were chosen and images were taken for the following 15-20 hours every 30 min. At the final time point, Draq 5 (1:1000) was added to each chamber and images were taken after 10min. Nikon Analysis software was used to analyse the relative distance between the borders of the wound and to visualize the nuclei in order to count the number of cells/area.

### Luciferase vectors construction and luciferase assay

To design the pGL3-CDCA2 wt plasmid, a 1367 bp region (CHR 8 25458872-25460239) of the CDCA2 promoter upstream the transcription start site (TSS, codon ATG) was cloned into the pGL3-Basic backbone vector upstream the luciferase gene. The pGL3-CDCA2 mut plasmid contains mutations of the canonical MYC binding sequence present in the wt segment (Fig 3D). Gene synthesis and cloning was performed by Biomatik Corporation.

For the luciferase assays, 5×10^4^/well of Ca1H cells were seeded in 24-well plates. The next day, cells were co-transfected with 0.25 μg of pGL3-CDCA2 (wt or mut) using JetPRIMER (Polyplus) and incubated for 36h. pRenilla luciferase vector was used to control transfection efficiency. Luciferase activity was detected with the Dual-Luciferase Reporter Assay System (Promega).

### Microarray analysis

MDA-MB-231 microarray analysis has been previously published (Masso-Valles *et al*, 2022). Briefly, MDA-MB-231 cells were seeded and on the next day were treated with 20 µmol/L Omomyc or with an equivalent volume of vehicle. After 3 days, plates were washed twice with PBS and frozen at−80°C until processing. RNA was extracted with TRIzol reagent (Invitrogen) according to the manufacturer’s instructions. The quality of RNA was confirmed with Agilent 2100 Bioanalyzer. Clariom S Human HT microarray plate (Applied Biosystems) was processed at Vall d’Hebron Institute of Research (VHIR)’s High Technology Unit. The microarray data were analyzed with Partek Genomics Suite software, v7.18. The expression heatmap was generated using R version 4.4.1 in R Studio version 2024.12.0+467, with the pheatmap package version 1.0.12 (Kolde R (2018). pheatmap: Pretty Heatmaps. R package version 1.0.12, https://github.com/raivokolde/pheatmap.).

### R2 Genomics analyses

The Genomics Analysis and Visualization Platform (https://hgserver1.amc.nl) was used to analyse the correlation of several RIPPOs genes across neuroblastoma datasets Kocak and SEQC or the Clynes dataset for mix breast tumors, and to generate the relative Kaplan Meier plots.

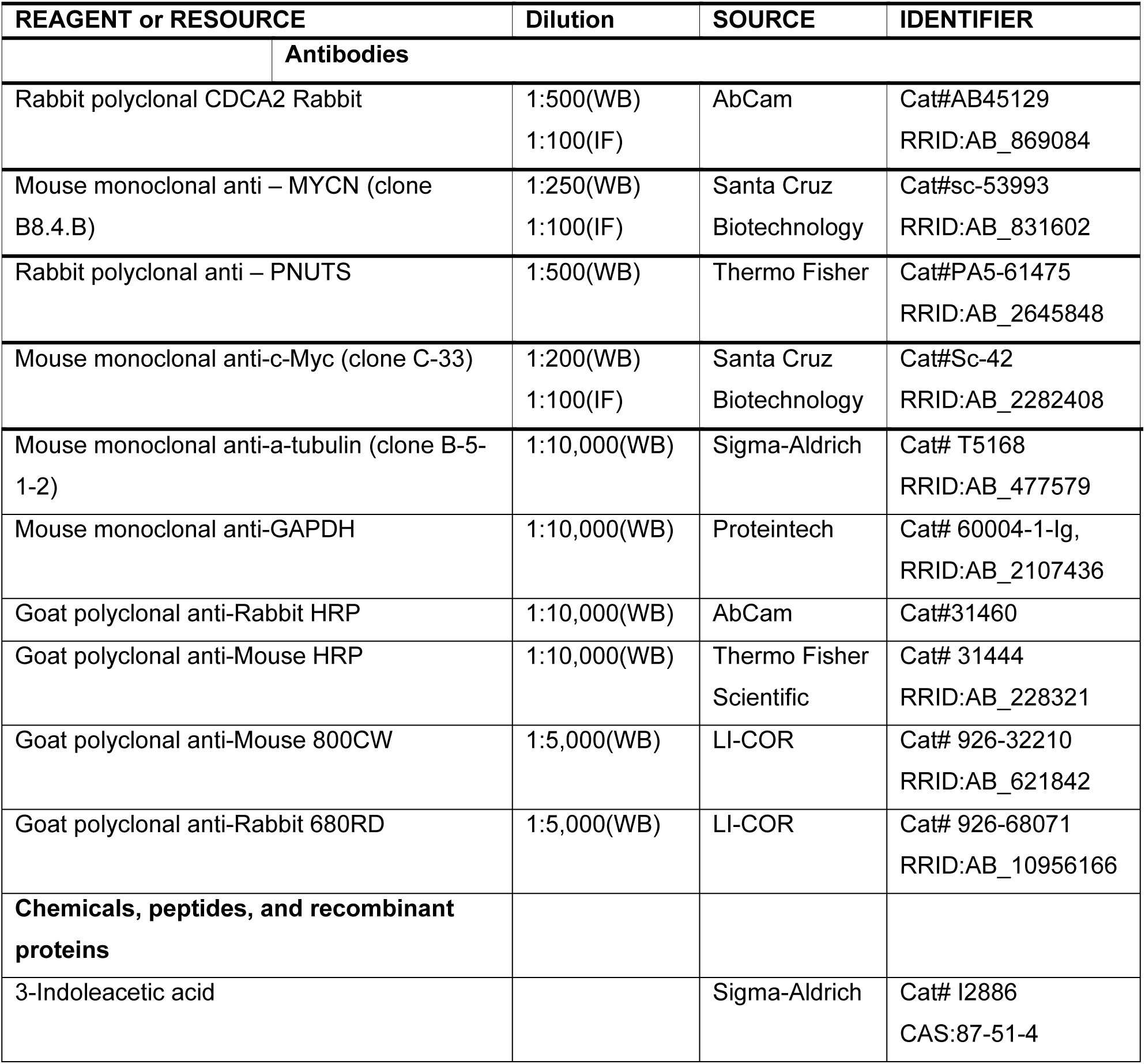
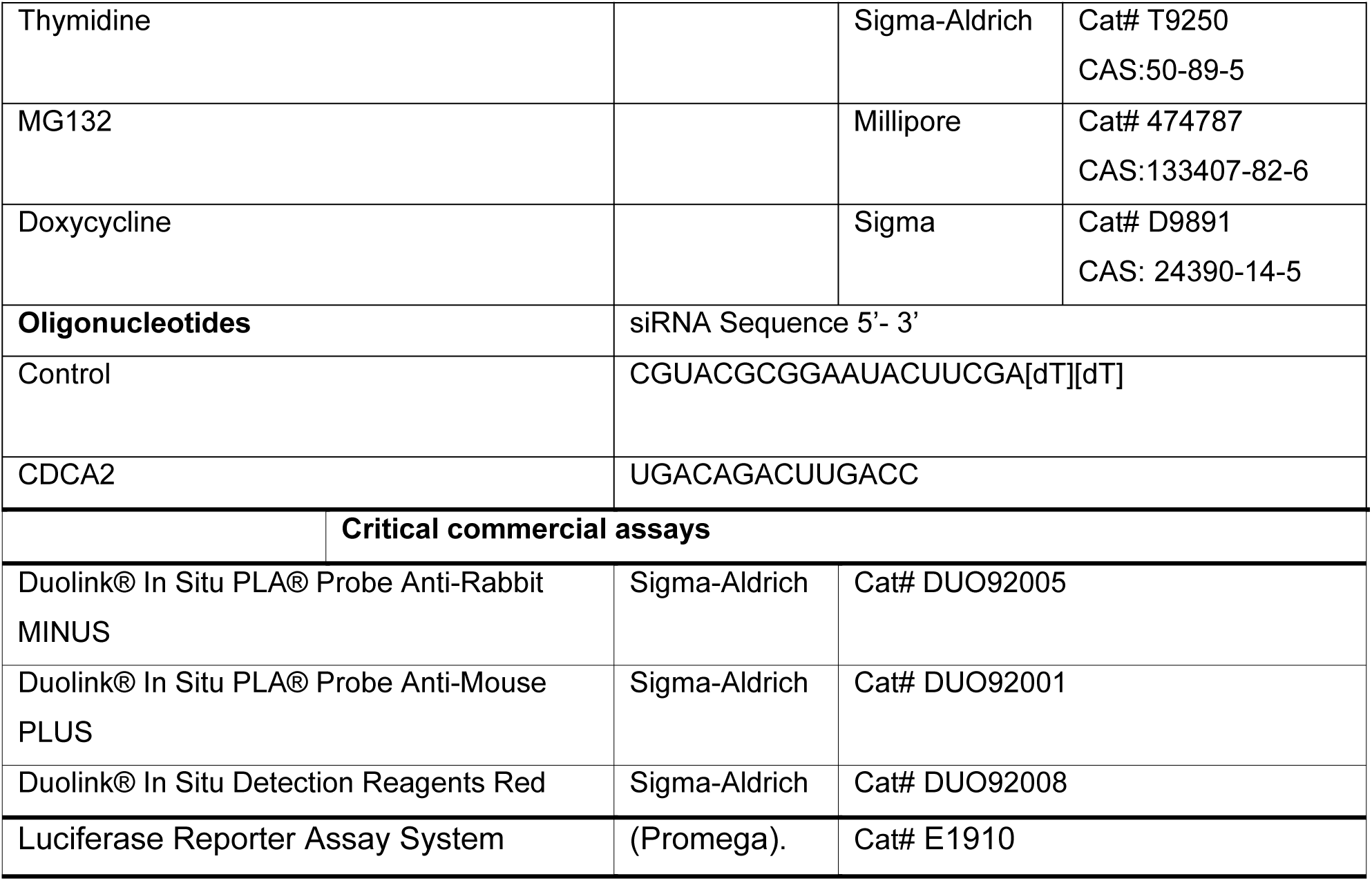

## Acknowledgements

The work was on the Breast Cancer was supported by a Breast Cancer Campaign PILOT grant to Paola Vagnarelli; The work on Neuroblastoma was supported by a KIDSCAN PhD studentship to PV and AS and a research grant from the Little Princess Trust to AS.

## Author Contribution

Conceptualisation: PV, AS; Methodology: KS, PV, LL, LS; Formal Analysis: KS, LL, MB, AT, EG CS, MFZ, AS, PV; Investigation: KS, LL, MB, AT, EG CS, MFZ, PV; Data Curation: KS, LS, MFZ, AS, PV; Writing – Review & Editing: PV and AS with contribution from all the authors; Visualization: PV, KS, AS; Supervision: PV, AS; Funding Acquisition: PV, AS

---

**Figure EV1.**
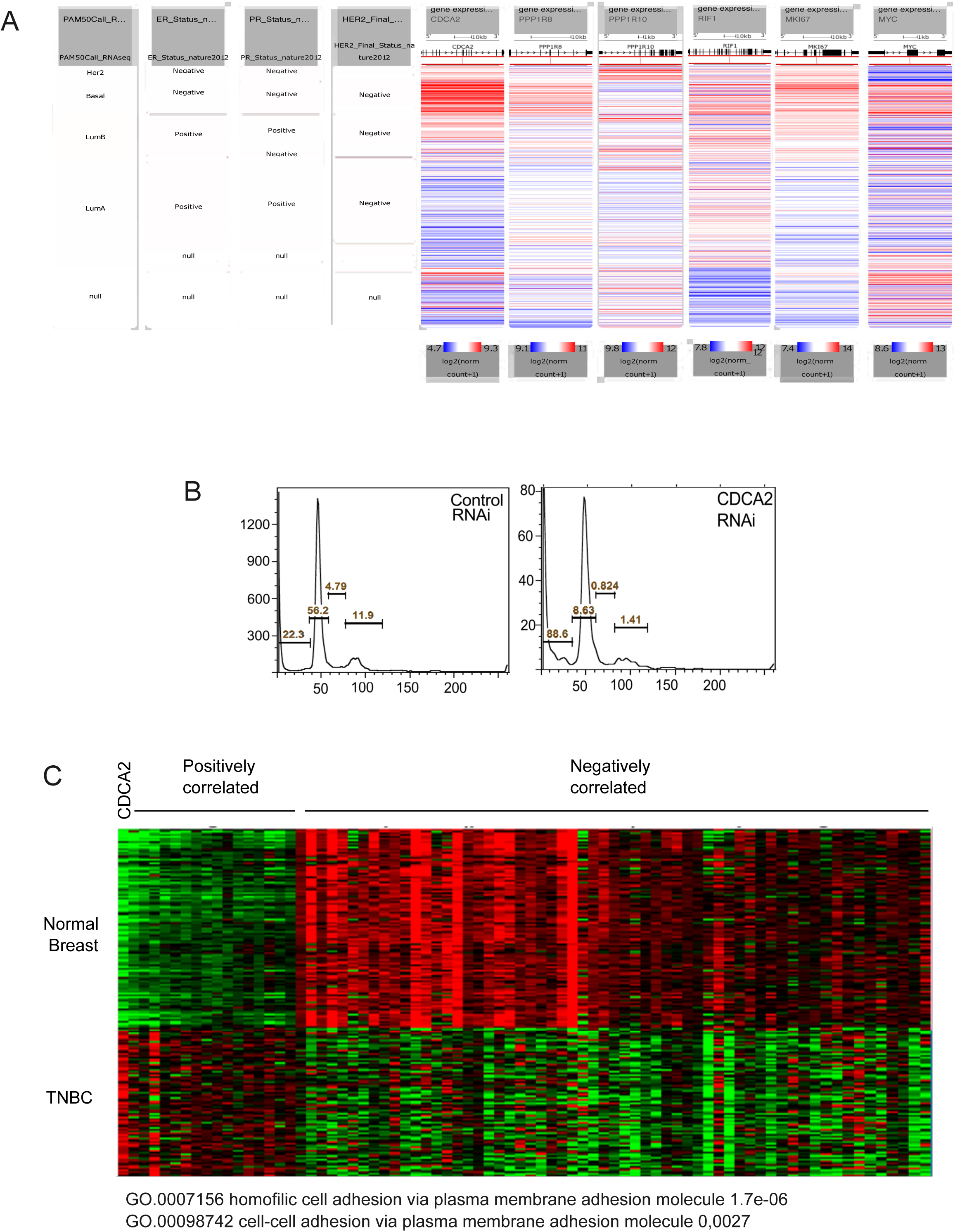
**A)** UCSC cancer browser analyses of CDCA2, PPP1R8, PPP1R10, RIF1and Ki-67 the expression levels compared to MYC expression in the TCGA breast database and stratified for the ER, PR and HER2 status. **B)** Flow cytometry profiles of Ca1H cell line transfected with control Si or CDCA2 Si oligos for 36 h. The numbers represent the percentage of cells in each stage: sub-G1, G1, S and G2/M. **C)** Expression levels of CDCA2 and genes bound by CDCA2 in normal breast and in TNBC tumour samples; in green is low expression and red high expression.

**Figure EV2.**
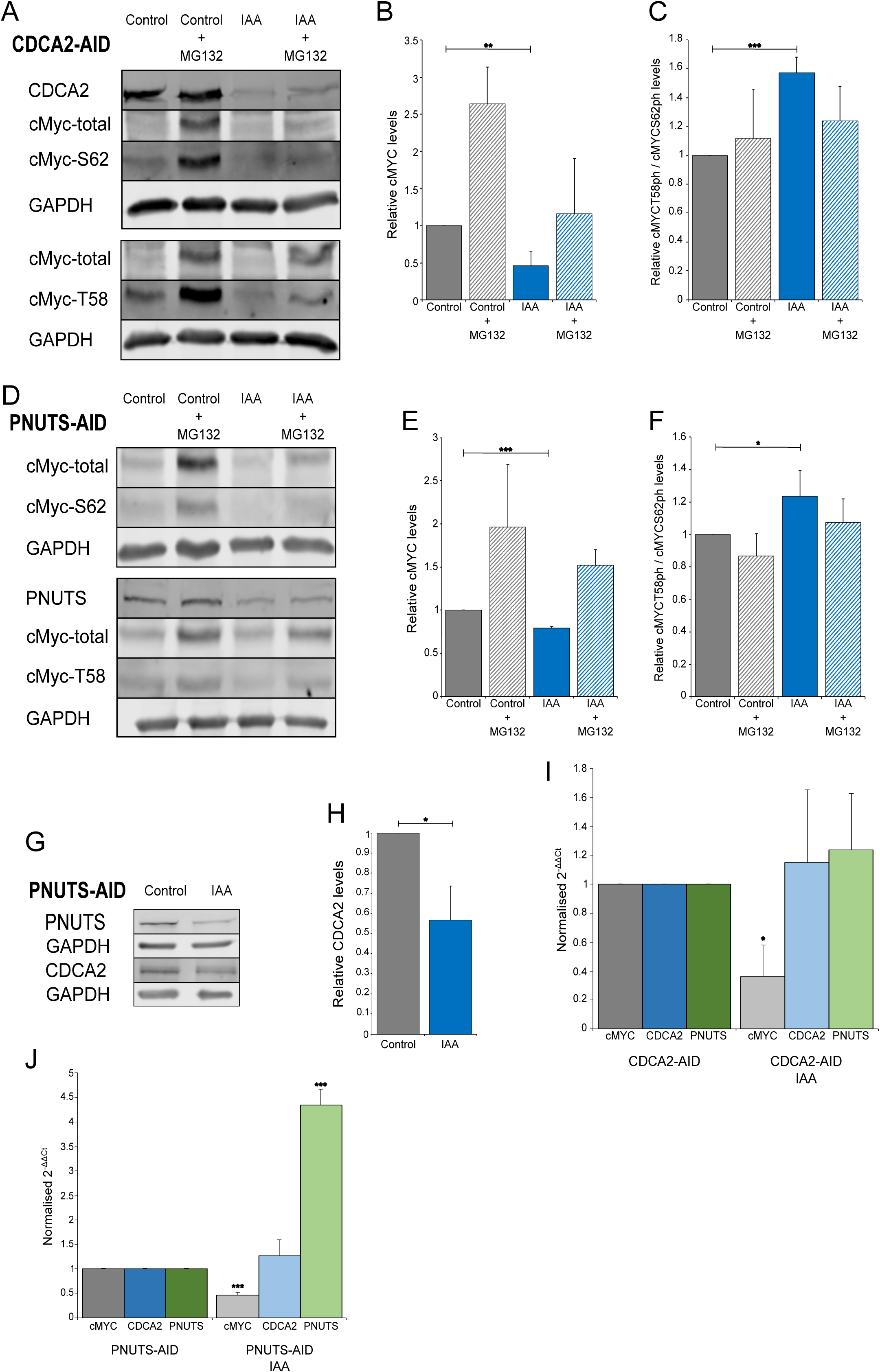
**A and D)** Representative Western blot analyses of HCT116-CDCA2-AID (A) and HCT116-PNUTS-AID (D) cell lines treated with doxycycline (2 μg/ml) 24 h and thymidine (2mM) for 18 h before the addition of IAA, then untreated (Control) or treated with IAA (IAA) for 4 h followed by 1 h treatment with 20μM MG132 in the presence (IAA + MG132) or absence (Control + MG132) of IAA. The blots were probed with anti-GAPDH, anti-CDCA2, anti-PNUTS, anti-cMYC-total, anti-cMYC-S62ph and anti-cMYC-T58ph antibodies. The images were acquired with a LICOR Imaging system in the linear range for quantification purposes. **B and E)** Quantification of total cMYC in HCT116-CDCA2-AID (A) and HCT116-PNUTS-AID (D) cell lines. The values represent the average of 3 independent replicas, and the error bars are the standard deviations. The experiments were analysed by Student’s t-test. **p < 0.01, ***p < 0.001. **C and F)** Quantification of cMYC-T58ph / cMYC-S62ph in HCT116-CDCA2-AID (A) and HCT116-PNUTS-AID (D) cell lines. The values represent the average of 3 independent replicas, and the error bars are the standard deviations. The experiments were analysed by Student’s t-test. ns=no significant, *p < 0.05, ***p < 0.001 **G)** Representative Western blot analyses of HCT116-PNUTS-AID cell line treated with doxycycline (2 μg/ml) for 24 h before the addition of IAA, then untreated (Control) or treated with IAA (IAA) for 4 h. **H)** Quantification of CDCA2 levels from the experiments in **(G)**. The values represent the average of 3 independent replicas, and the error bars are the standard deviations. The experiments were analysed by Student’s t-test. ns=no significant, *p < 0.05. **I and J)** qPCR analyses of cMYC, CDCA2 and PNUTS expression in HCT116-CDCA2-AID (I) and HCT116-PNUTS-AID (J) cell lines, treated with doxycycline (2 μg/ml) 24 h and thymidine (2mM) for 18 h before the addition of IAA for 4 h. The data represent the average of 3 independent replicas and the error bars are the standard deviations. The experiments were analysed by Student’s t-test. *p < 0.05, ***p < 0.001

**Figure EV3.**
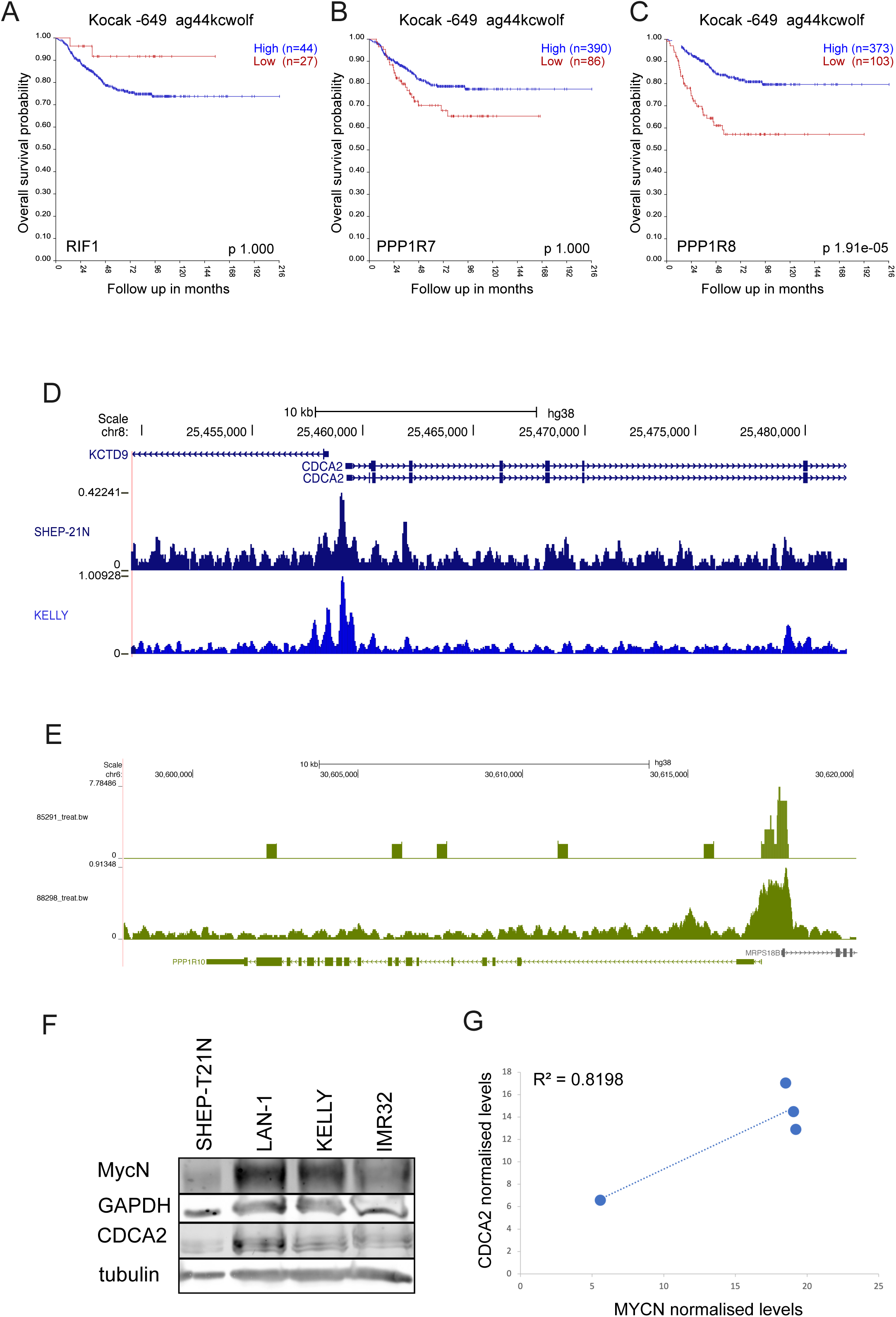
**A)** Kaplan–Meier curve (survival probability in months) of neuroblastoma patients expressing low (red curve) or high (blue curve) levels of RIF1 (Kocak cohort). **B)** Kaplan–Meier curve (survival probability in months) of neuroblastoma patients expressing low (red curve) or high (blue curve) levels of PPP1R7 (Kocak cohort). **C)** Kaplan–Meier curve (survival probability in months) of neuroblastoma patients expressing low (red curve) or high (blue curve) levels of PPP1R8 (Kocak cohort). **D)** MYCN chromatin immunoprecipitation sequencing profiles at the PNUTS locus in Kelly cell lines. **E)** MYCN chromatin immunoprecipitation sequencing profiles at the CDCA2 locus in SHEP-T21N and KELLY cell lines. **F)** Western blot analyses of SHEP-T21N, LAN-1, KELLY and IMR32 cell lines. The blots were probed with anti-alpha tubulin or anti-GAPDH and with anti-CDCA2, anti-MYCN-total antibodies. The images were acquired with a LICOR Imaging system in the linear range for quantification purposes. **G)** Correlation of MYCN to CDCA2 normalised protein levels from (F).

**Figure EV4.**
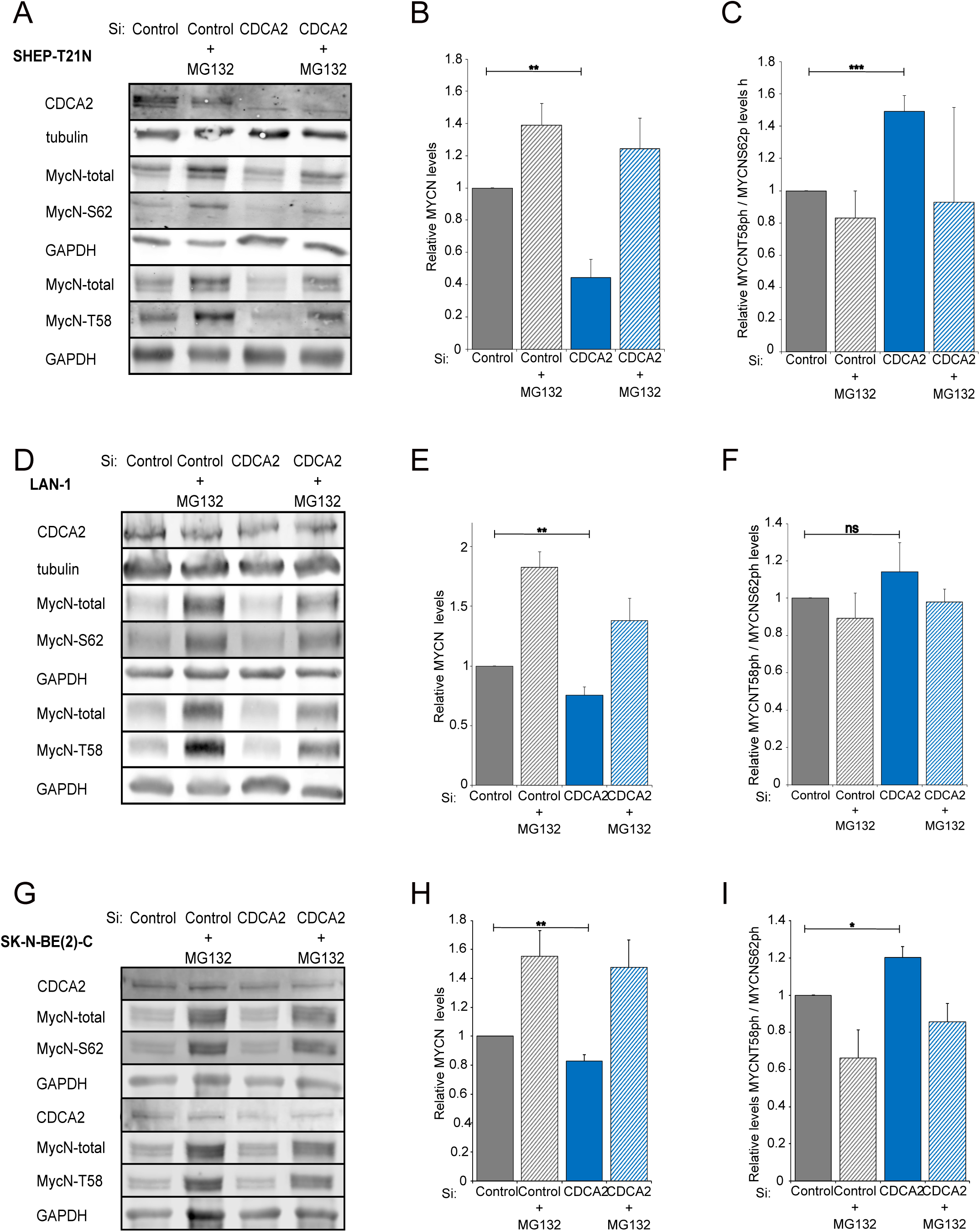
**A, D and G)** Representative Western blot analyses of SHEP-T21N (A), LAN-1 (D) and SK-N-BE(2)-C cell lines. 48 h post transfection with control or CDCA2 Si oligos, cells were treated with 20μM MG132 for 2 h. The blots were probed with anti-alpha tubulin or anti-GAPDH and with anti-CDCA2, anti-MYCN-total, anti-cMYC-S62ph and anti-cMYC-T58ph antibodies. The images were acquired with a LICOR Imaging system in the linear range for quantification purposes. **B, E and H)** Quantification of total MYCN in SHEP-T21N (A), LAN-1 (D) and SK-N-BE(2)-C cell lines. The values represent the average of 3 independent replicas, and the error bars are the standard deviations. The experiments were analysed by Student’s t-test. **p < 0.01. **C, F and I)** Quantification of MYCN-T58ph / MYCN-S62ph in SHEP-T21N (A), LAN-1 (D) and SK-N-BE(2)-C cell lines. The values represent the average of 3 independent replicas, and the error bars are the standard deviations. The experiments were analysed by Student’s t-test. ns=no significant, *p < 0.05, ***p < 0.001

## REFERENCES

Baluapuri A, Wolf E, Eilers M (2020) Target gene-independent functions of MYC oncoproteins. Nat Rev Mol Cell Biol 21: 255–267

Baxter JS, Brough R, Krastev DB, Song F, Sridhar S, Gulati A, Alexander J, Roumeliotis TI, Kozik Z, Choudhary JS et al (2024) Cancer-associated FBXW7 loss is synthetic lethal with pharmacological targeting of CDC7. Mol Oncol 18: 369–385

Boi D, Rubini E, Breccia S, Guarguaglini G, Paiardini A (2023) When Just One Phosphate Is One Too Many: The Multifaceted Interplay between Myc and Kinases. Int J Mol Sci 24

Corvetta D, Chayka O, Gherardi S, D’Acunto CW, Cantilena S, Valli E, Piotrowska I, Perini G, Sala A (2013) Physical interaction between MYCN oncogene and polycomb repressive complex 2 (PRC2) in neuroblastoma: functional and therapeutic implications. J Biol Chem 288: 8332–8341

de Castro IJ, Budzak J, Di Giacinto ML, Ligammari L, Gokhan E, Spanos C, Moralli D, Richardson C, de Las Heras JI, Salatino S et al (2017) Repo-Man/PP1 regulates heterochromatin formation in interphase. Nat Commun 8: 14048

Dingar D, Tu WB, Resetca D, Lourenco C, Tamachi A, De Melo J, Houlahan KE, Kalkat M, Chan PK, Boutros PC et al (2018) MYC dephosphorylation by the PP1/PNUTS phosphatase complex regulates chromatin binding and protein stability. Nat Commun 9: 3502

Diolaiti D, McFerrin L, Carroll PA, Eisenman RN (2015) Functional interactions among members of the MAX and MLX transcriptional network during oncogenesis. Biochim Biophys Acta 1849: 484–500

Egger JV, Lane MV, Antonucci LA, Dedi B, Krucher NA (2016) Dephosphorylation of the Retinoblastoma protein (Rb) inhibits cancer cell EMT via Zeb. Cancer Biol Ther 17: 1197–1205

Eilers M, Eisenman RN (2008) Myc’s broad reach. Genes Dev 22: 2755–2766

Fallah Y, Brundage J, Allegakoen P, Shajahan-Haq AN (2017) MYC-Driven Pathways in Breast Cancer Subtypes. Biomolecules 7

Finkelman BS, Zhang H, Hicks DG, Turner BM (2023) The Evolution of Ki-67 and Breast Carcinoma: Past Observations, Present Directions, and Future Considerations. Cancers (Basel) 15

Garralda E, Beaulieu ME, Moreno V, Casacuberta-Serra S, Martinez-Martin S, Foradada L, Alonso G, Masso-Valles D, Lopez-Estevez S, Jauset T et al (2024) MYC targeting by OMO-103 in solid tumors: a phase 1 trial. Nat Med 30: 762–771

Gaubatz S, Meichle A, Eilers M (1994) An E-box element localized in the first intron mediates regulation of the prothymosin alpha gene by c-myc. Mol Cell Biol 14: 3853–3862

Gu P, Yang D, Zhu J, Zhang M, He X (2022) Bioinformatics analysis of the clinical relevance of CDCA gene family in prostate cancer. Medicine (Baltimore) 101: e28788

Harvey-Jones E, Raghunandan M, Robbez-Masson L, Magraner-Pardo L, Alaguthurai T, Yablonovitch A, Yen J, Xiao H, Brough R, Frankum J et al (2024) Longitudinal profiling identifies co-occurring BRCA1/2 reversions, TP53BP1, RIF1 and PAXIP1 mutations in PARP inhibitor-resistant advanced breast cancer. Ann Oncol 35: 364–380

Hydbring P, Bahram F, Su Y, Tronnersjo S, Hogstrand K, von der Lehr N, Sharifi HR, Lilischkis R, Hein N, Wu S et al (2010) Phosphorylation by Cdk2 is required for Myc to repress Ras-induced senescence in cotransformation. Proc Natl Acad Sci U S A 107: 58–63

Hydbring P, Larsson LG (2010) Tipping the balance: Cdk2 enables Myc to suppress senescence. Cancer Res 70: 6687–6691

Jakobsen ST, Siersbaek R (2025) Transcriptional regulation by MYC: an emerging new model. Oncogene 44: 1–7

Jiang J, Wang J, Yue M, Cai X, Wang T, Wu C, Su H, Wang Y, Han M, Zhang Y et al (2020) Direct Phosphorylation and Stabilization of MYC by Aurora B Kinase Promote T-cell Leukemogenesis. Cancer Cell 37: 200–215 e205

Kalkat M, Resetca D, Lourenco C, Chan PK, Wei Y, Shiah YJ, Vitkin N, Tong Y, Sunnerhagen M, Done SJ et al (2018) MYC Protein Interactome Profiling Reveals Functionally Distinct Regions that Cooperate to Drive Tumorigenesis. Mol Cell 72: 836–848 e837

Kim JY, Valencia T, Abu-Baker S, Linares J, Lee SJ, Yajima T, Chen J, Eroshkin A, Castilla EA, Brill LM et al (2013) c-Myc phosphorylation by PKCzeta represses prostate tumorigenesis. Proc Natl Acad Sci U S A 110: 6418–6423

Krasnoselsky AL, Whiteford CC, Wei JS, Bilke S, Westermann F, Chen QR, Khan J (2005) Altered expression of cell cycle genes distinguishes aggressive neuroblastoma. Oncogene 24: 1533–1541

Laemmli UK (1970) Cleavage of structural proteins during the assembly of the head of bacteriophage T4. Nature 227: 680–685

Lashen AG, Toss MS, Ghannam SF, Makhlouf S, Green A, Mongan NP, Rakha E (2023) Expression, assessment and significance of Ki67 expression in breast cancer: an update. J Clin Pathol 76: 357–364

Li W, Lv D, Yao J, Chen B, Liu H, Li W, Xu C, Li Z (2023) A pan-cancer analysis reveals the diagnostic and prognostic role of CDCA2 in low-grade glioma. PLoS One 18: e0291024

Lu JJ, Meng LH, Shankavaram UT, Zhu CH, Tong LJ, Chen G, Lin LP, Weinstein JN, Ding J (2010) Dihydroartemisinin accelerates c-MYC oncoprotein degradation and induces apoptosis in c-MYC-overexpressing tumor cells. Biochem Pharmacol 80: 22–30

Lutz W, Stohr M, Schurmann J, Wenzel A, Lohr A, Schwab M (1996) Conditional expression of N-myc in human neuroblastoma cells increases expression of alpha-prothymosin and ornithine decarboxylase and accelerates progression into S-phase early after mitogenic stimulation of quiescent cells. Oncogene 13: 803–812

Manzione MG, Rombouts J, Steklov M, Pasquali L, Sablina A, Gelens L, Qian J, Bollen M (2020) Co-regulation of the antagonistic RepoMan:Aurora-B pair in proliferating cells. Mol Biol Cell 31: 419–438

Marx A, Luebke AM, Clauditz TS, Steurer S, Fraune C, Hube-Magg C, Buscheck F, Hoflmayer D, Tsourlakis MC, Moller-Koop C et al (2020) Upregulation of Phosphatase 1 Nuclear-Targeting Subunit (PNUTS) Is an Independent Predictor of Poor Prognosis in Prostate Cancer. Dis Markers 2020: 7050146

Masso-Valles D, Beaulieu ME, Jauset T, Giuntini F, Zacarias-Fluck MF, Foradada L, Martinez-Martin S, Serrano E, Martin-Fernandez G, Casacuberta-Serra S et al (2022) MYC Inhibition Halts Metastatic Breast Cancer Progression by Blocking Growth, Invasion, and Seeding. Cancer Res Commun 2: 110–130

Masso-Valles D, Soucek L (2020) Blocking Myc to Treat Cancer: Reflecting on Two Decades of Omomyc. Cells 9

Myant K, Qiao X, Halonen T, Come C, Laine A, Janghorban M, Partanen JI, Cassidy J, Ogg EL, Cammareri P et al (2015) Serine 62-Phosphorylated MYC Associates with Nuclear Lamins and Its Regulation by CIP2A Is Essential for Regenerative Proliferation. Cell Rep 12: 1019–1031

Peng A, Lewellyn AL, Schiemann WP, Maller JL (2010) Repo-man controls a protein phosphatase 1-dependent threshold for DNA damage checkpoint activation. Curr Biol 20: 387–396

Qian J, Beullens M, Lesage B, Bollen M (2013) Aurora B defines its own chromosomal targeting by opposing the recruitment of the phosphatase scaffold Repo-Man. Curr Biol 23: 1136–1143

Qian J, Lesage B, Beullens M, Van Eynde A, Bollen M (2011) PP1/Repo-man dephosphorylates mitotic histone H3 at T3 and regulates chromosomal aurora B targeting. Curr Biol 21: 766–773

Sabo A, Amati B (2014) Genome recognition by MYC. Cold Spring Harb Perspect Med 4

Solvie D, Baluapuri A, Uhl L, Fleischhauer D, Endres T, Papadopoulos D, Aziba A, Gaballa A, Mikicic I, Isaakova E et al (2022) MYC multimers shield stalled replication forks from RNA polymerase. Nature 612: 148–155

Stamatiou K, Huguet F, Budzinska M, Ragusa D, deCastro I, J,, Spanos C, Rappsilberg J, Vagnarelli P (2024) Multi-omic analyses reveal a differential contribution of chromatin-associated PP1 holoenzymes to mitotic exit and G1 re-establishment. Bioarchive

Sun W, Jin Y, Wei C, Xu Y, Liu W, Zhong J, Zou Z, Lin X, Xiang Y, Chen Y (2023) CDCA2 promotes melanoma progression by inhibiting ubiquitin-mediated degradation of Aurora kinase A. Eur J Cancer 188: 49–63

Tang M, Liao M, Ai X, He G (2021) Increased CDCA2 Level Was Related to Poor Prognosis in Hepatocellular Carcinoma and Associated With Up-Regulation of Immune Checkpoints. Front Med (Lausanne) 8: 773724

Trinkle-Mulcahy L, Andersen J, Lam YW, Moorhead G, Mann M, Lamond AI (2006) Repo-Man recruits PP1 gamma to chromatin and is essential for cell viability. J Cell Biol 172: 679–692

Uchida F, Uzawa K, Kasamatsu A, Takatori H, Sakamoto Y, Ogawara K, Shiiba M, Bukawa H, Tanzawa H (2013) Overexpression of CDCA2 in human squamous cell carcinoma: correlation with prevention of G1 phase arrest and apoptosis. PLoS One 8: e56381

Vagnarelli P (2014) Repo-man at the intersection of chromatin remodelling, DNA repair, nuclear envelope organization, and cancer progression. Adv Exp Med Biol 773: 401–414

Vagnarelli P, Hudson DF, Ribeiro SA, Trinkle-Mulcahy L, Spence JM, Lai F, Farr CJ, Lamond AI, Earnshaw WC (2006) Condensin and Repo-Man-PP1 co-operate in the regulation of chromosome architecture during mitosis. Nat Cell Biol 8: 1133–1142

Vagnarelli P, Ribeiro S, Sennels L, Sanchez-Pulido L, de Lima Alves F, Verheyen T, Kelly DA, Ponting CP, Rappsilber J, Earnshaw WC (2011) Repo-Man coordinates chromosomal reorganization with nuclear envelope reassembly during mitotic exit. Dev Cell 21: 328–342

van de Wetering M, Sancho E, Verweij C, de Lau W, Oving I, Hurlstone A, van der Horn K, Batlle E, Coudreuse D, Haramis AP et al (2002) The beta-catenin/TCF-4 complex imposes a crypt progenitor phenotype on colorectal cancer cells. Cell 111: 241–250

Wang L, Chen C, Song Z, Wang H, Ye M, Wang D, Kang W, Liu H, Qing G (2022) EZH2 depletion potentiates MYC degradation inhibiting neuroblastoma and small cell carcinoma tumor formation. Nat Commun 13: 12

Wei Y, Redel C, Ahlner A, Lemak A, Johansson-Akhe I, Houliston S, Kenney TMG, Tamachi A, Morad V, Duan S et al (2022) The MYC oncoprotein directly interacts with its chromatin cofactor PNUTS to recruit PP1 phosphatase. Nucleic Acids Res 50: 3505–3522

Zacarias-Fluck MF, Soucek L, Whitfield JR (2024) MYC: there is more to it than cancer. Front Cell Dev Biol 12: 1342872

Zhou Y, Chen W, Jiang H, Zhang Y, Ma Z, Wang Z, Xu C, Jiang M, Chen J, Cao Z (2024) MKI67 with arterial hypertension predict a poor survival for prostate cancer patients, a real-life investigation. Clin Transl Oncol 26: 3037–3049

